# Single-cell Mendelian randomisation identifies cell-type specific genetic effects on human brain disease and behaviour

**DOI:** 10.1101/2022.11.28.517913

**Authors:** Alexander Haglund, Verena Zuber, Yifei Yang, Maya Abouzeid, Rahel Feleke, Jeong Hun Ko, Alexi Nott, Ann C. Babtie, James D. Mills, Louwai Muhammed, Liisi Laaniste, Djordje O. Gveric, Daniel Clode, Susanna Pagni, Ravishankara Bellampalli, Alyma Somani, Karina McDade, Jasper J. Anink, Lucia Mesarosova, Eleonora Aronica, Maria Thom, Sanjay M. Sisodiya, Prashant K. Srivastava, Dheeraj Malhotra, Julien Bryois, Leonardo Bottolo, Michael R. Johnson

**Author notes:** Authors contributed equally to this work (Alexander Haglund, Verena Zuber). Senior author (Maria Thom, Sanjay M. Sisodiya, Prashant K. Srivastava, Dheeraj Malhotra, Julien Bryois, Leonardo Bottolo, Michael R. Johnson).

## Abstract

Translating genome-wide association loci to therapies requires knowledge of the causal genes, their directionality of effect and the cell-types in which they act. To infer these relationships in the human brain, we implemented Mendelian randomisation using single cell-type expression quantitative trait loci (eQTLs) as genetic anchors. Expression QTLs were mapped across 8 major cell-types in brain tissue exclusively ascertained from donors with no history of brain disease. We report evidence for a causal association between the change in expression of 118 genes and one or more of 16 brain phenotypes, revealing candidate targets for risk mitigation and opportunities for shared preventative therapeutic strategies. We highlight key causal genes for neurodegenerative and neuropsychiatric disease and for each, we report its cellular context and the therapeutic directionality required for risk mitigation. Our use of control samples establishes a new resource for the causal interpretation of GWAS risk alleles for human brain phenotypes.

## INTRODUCTION

The average cost to bring a drug to market is $2.6 billion (2013 dollars)[1]. Only 4% of drug-development programs yield licensed drugs due to two main issues: (a) preclinical experimental models are poorly predictive of eventual therapeutic efficacy and (b) definitive evidence of target validity is not obtained until randomised controlled trials (RCT) in late-stage drug development[2]. The retrospective observation that drugs with genetic support for the target-indication pairing are more than twice as likely to be successful in clinical development has therefore focused attention on the potential for human genetics to predict successful new drugs[3], [4]. However, translating genetic loci to therapies requires knowledge of the causal genes as well as the directionality of effect of a gene’s expression on disease risk in specific cell-types, which is rarely directly available from genetic analysis alone[5],[6].

Here, we aimed to infer these causal relationships by implementing a principled approach to Mendelian randomisation (MR) using single cell-type expression quantitative trait loci (eQTLs) as genetic anchors. MR is a statistical framework for inferring causal associations using human observational data[7]. Instead of randomising subjects to drug exposure versus placebo to investigate the causal relation between an exposure and a health outcome, MR makes use of the naturally randomized allocation of genetic variants (SNPs) that instrument an exposure such as the level of expression of a gene[8].

In the present study, we restricted our analysis to human brain single-cell gene expression data ascertained exclusively from donors with no history of brain disease and with normal appearances of the brain on neuropathological examination. Although brain tissue samples from people who have died with a neurological or psychiatric diagnosis are more widely available than control samples, the use of diseased brain tissue has the potential to confound the deconvolution of true forward causal effects from mere correlation due to biased anchoring of the causal inference in disease-induced gene expression changes rather than disease-causing ones (confounding by reverse causation)[9]. In contrast, our use of brain tissue that predates the onset of brain disease offers an opportunity to discover cell-type specific causal risk factors that are unconfounded by reverse causation and therefore modifiable drug targets for disease prevention. By focussing solely on control samples, we establish a new resource for the interpretation of GWAS-risk alleles on human brain phenotypes.

In addition to providing an improved level of certainty about the causal relation between a candidate drug target and a clinical outcome, the application of MR anchored in single cell-type eQTLs also provides estimates of the size and direction of the effect of an exposure on an outcome in a specific cell-type. These estimates are critical to designing the correct therapeutic intervention. Therefore, to enable a transparent assessment of our cell-type specific causal inferences we report our findings in line with the STROBE-MR guidelines for MR studies[10], including explicit reporting of the strength of the statistical evidence at each step.

## RESULTS

### Study overview

To study cell-type specific genetic effects on human brain structure, disease, and behaviour we utilized single-nuclei gene expression data (snRNA-seq) based on post-mortem brain tissue samples from 147 genotyped adult donors. Across all donors, there was no history of neurological or psychiatric disease prior to death, and no evidence for disease of the brain on neuropathological examination. Single cell-type Mendelian randomisation (MR) analysis was implemented on this resource in three stages: (a) data generation and single cell-type eQTL mapping, (b) instrumental variable selection and assessment, (c) two-sample Mendelian randomisation (MR) (study design summarised in Fig.1a).

**Fig. 1.**
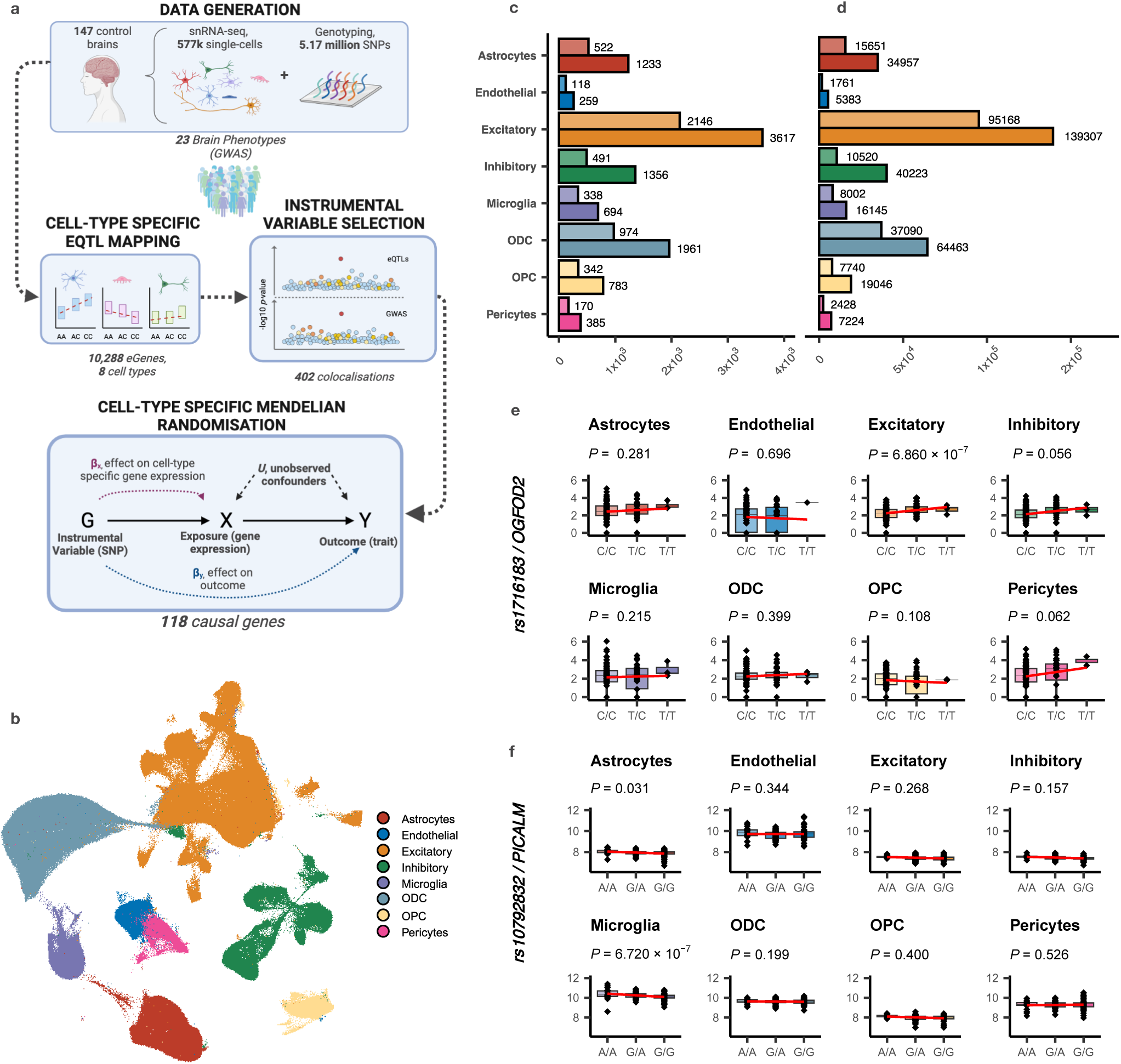
Study overview, cell types and single cell-type *cis*-eQTLs. **a**. Single cell-type *cis*-eQTLs were identified in control brain tissue samples and integrated with GWAS loci in a Mendelian randomisation framework to infer the causal relationships between genes, cell-types and phenotypes. **b**. The eight major cell-types of the human brain (excitatory neurons, oligodendrocytes, astrocytes, inhibitory neurons, microglia, oligodendrocyte precursor cells, endothelial cells and pericytes) were identified from snRNA-seq using canonical cell-type markers. **c**. Number of eGenes unique (top line) and total (bottom line) for each cell-type at <5% FDR. **d**. Number of *cis*-eQTLs eSNPs unique (top line) and total (bottom line) for each cell-type at <5% FDR. **e**. An example of a cell-type-specific *cis*-eQTL (SNP-gene pair) in excitatory neurons. **f**. An example of cell-type specific *cis*-eQTL (SNP-gene pair) in microglia.

### Data generation and single cell-type *cis-*eQTL mapping

After quality control, sample integration, cell-type annotation and genotype imputation, 577,115 single-cells across 128 subjects averaging 4,509 cells per donor were available for estimating allele-specific effects on gene expression in single cell-types (hereon referred to as single cell-type eQTLs). The 577,115 single cells across the sample set were aligned in a single graph (Fig. 1b) and consisted of 219,942 excitatory neurons, 66,246 inhibitory neurons, 133,752 oligodendrocytes, 68,809 astrocytes, 30,086 microglia, 27,248 oligodendrocyte precursors, 17,144 endothelial cells and 13,888 pericytes (overview of snRNA-seq data characteristics in Supplementary Fig.1).

To calculate cell-type specific eQTLs we generated pseudobulk gene expression matrices by aggregating read counts for each gene in each cell-type for each subject (Methods). *cis-*eQTLs were mapped using MatrixEQTL[11] for each SNP-gene pair in each cell-type using a *cis* window extending 1Mb either side of the gene per protocol and adjusting for age, sex, post-mortem interval, sample source and the first 40 principal components of gene expression as fixed covariates[12]. In total, across the eight cell-types, 326,748 *cis-*eQTLs were identified at a study-wide False Discovery Rate (FDR) <5% [13] corresponding to one or more regulatory SNP (eSNP) for 10,288 genes (eGenes) (Figs.1c-d). Of these, 5,101 eGenes were unique to a single cell-type (illustrative examples in Figs.1e-f). Across the set of single cell-type *cis-*eQTLs, we observed a high level of replication (71.3-83.6%, varying by cell-type) in a large independent *cis-*eQTL dataset derived from bulk brain tissue samples from 6,518 subjects [14](Supplementary Figure 2).

### Instrumental variable selection

Valid genetic instruments for MR are underpinned by three core assumptions: They are associated with the exposure of interest (the relevance assumption); they only act via the measured exposure (the exclusion restriction assumption); there are no unmeasured confounders of the association between the genetic instrument and the outcome (the independence assumption)[15].

To plausibly meet these assumptions, we took a principled approach to the selection of instrumental variables (IVs). As a first step, we assessed whether phenotypic outcomes and potential gene mediators might share one or more causal variants using colocalization analysis. COLOC[16] is a method for genetic colocalization analysis that provides an estimate of the posterior probability of a shared signal between pairs of genetic association studies – in our case between a cell-type specific *cis-*eQTL (i.e., a SNP-gene pair in a particular cell-type) as one “trait”, and a SNP-phenotype association from a well-conducted GWAS as the second. We restricted the colocalization analysis to chromosomal regions containing a genome-wide significant association with the outcome in question (defined as a GWAS *P* <5.0×10^−8^). Colocalization analysis was carried out across 23 human brain phenotypes and the resulting cell-type specific colocalizations are summarised for each outcome in Supplementary Fig.3. As an illustrative example, we show the cell-type specific posterior probability of colocalizations (PP.H4>0.5) with Alzheimer’s disease (AD) in Fig.2a. These reveal several genes in specific cell-types concordant with the known biology of AD such as *PICALM* (PP.H4 microglia = 0.99; Figs.2b-c) and *RIN3* (PP.H4 microglia = 0.99)[17], as well as genes with a previously proposed but less well-established link to AD such as *SNX31* (PP.H4 astrocytes = 0.99)[18]. In total, across all phenotypes, we identified 402 cell-type specific colocalizations with PP.H4>0.5 (summary of the number of colocalised genes and cell-types for each brain phenotype in Fig.2d and Fig.2e respectively).

**Fig. 2.**
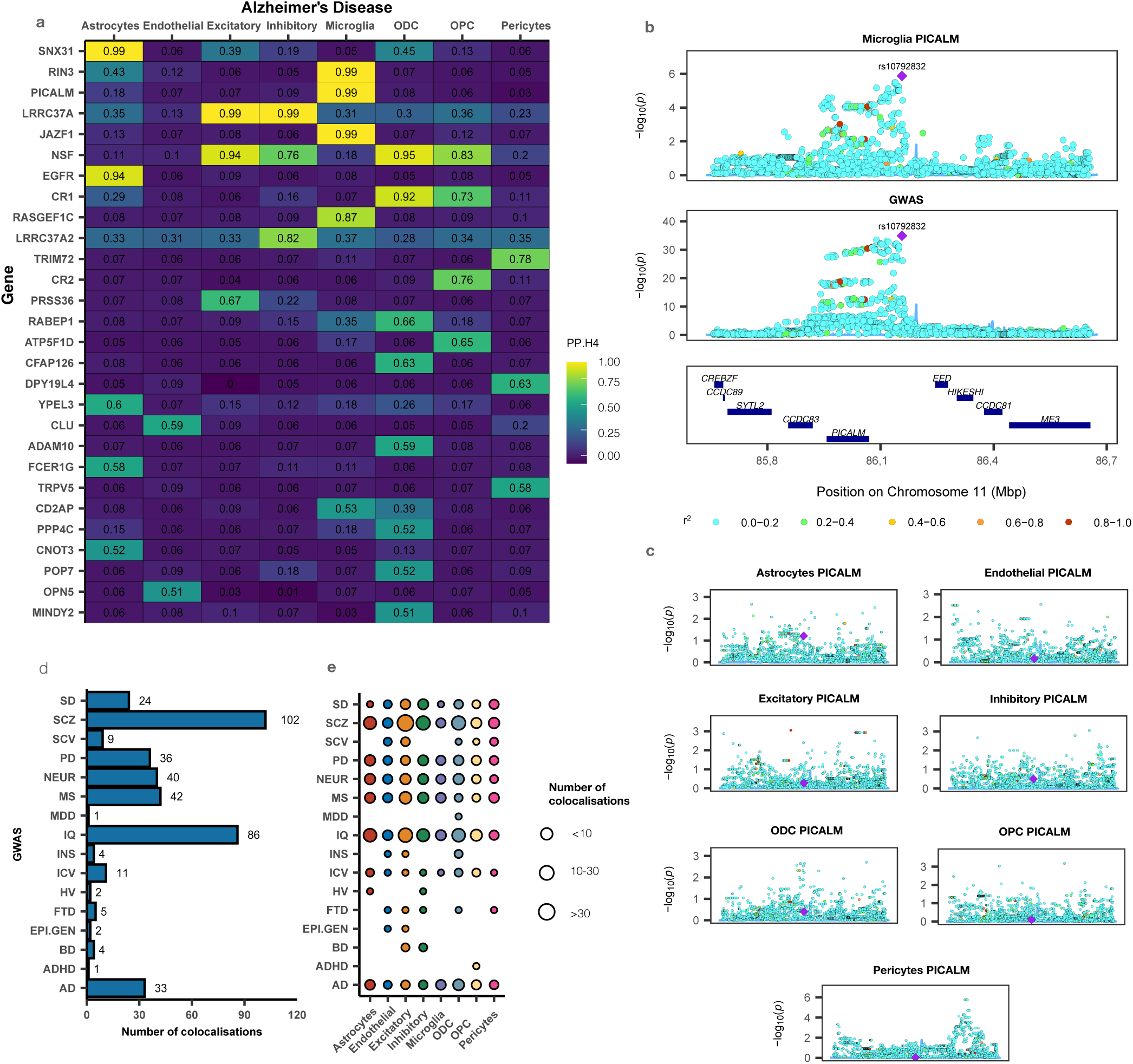
Colocalization analysis. **a**. Heatmap of posterior probability (PP.H4>0.5) for a shared genetic signal for a SNP-gene (i.e., *cis*-eQTL) pair (row) in a particular cell-type (column) and a genome-wide significant GWAS locus for Alzheimer’s disease (AD). **b**. Example of a microglial-specific colocalization between PICALM *cis*-eQTLs and AD. Each blue circle represents a SNP with the significance of its association (*y*-axis) to PICALM expression (top) or AD (bottom). SNP rs10792832 (purple diamond) is the lead colocalised SNP across the two associations. **c**. SNP-PICALM associations in the other cell-types across the same chromosomal region illustrating the lack of colocalization in other cell-types. **d**. Summary of the number of colocalizations (PP.H4>0.5) for each phenotype (SD: sleep duration; SCZ: schizophrenia; SCV: subcortical volume caudate; PD: Parkinson’s disease; NEUR: neuroticism; MS: multiple sclerosis; MDD: major depressive disorder; IQ: intelligence; INS: insomnia; ICV: intracranial volume; HV: hippocampal volume; FTD: Frontotemporal Dementia; EPI.GEN: genetic generalized epilepsy; BD: bipolar disorder; ADHD: attention deficit hyperactivity disorder, AD: Alzheimer disease). Each cell-type/gene pair with PP.H4>0.5 is reported - for example, LRRC37A has two colocalisations with AD, one in Excitatory Neurons and one in Inhibitory Neurons and therefore counts for two colocalisations. **e**. Bubble plot demonstrating the number of occurrences of a particular cell-type in a colocalization for the indicated phenotype.

To select the specific IV SNPs for MR analysis we first retained only the colocalised regions with a posterior probability (PP.H4) >0.5 for a shared causal signal, of which 76.5% mapped to a single cell-type. In line with the relevance assumption, we removed all SNPs in the colocalised region with a study-wide *cis*-eQTL FDR>5%. We then identified the lead eQTL SNP in the colocalised region and removed all variants in linkage disequilibrium (LD r^2^>0.01) with that SNP so as to minimise the risk of confounding by LD (i.e., confounding because the genetic variant is in LD with another variant that independently influences the outcome via an alternative unmeasured risk factor). For the retained SNPs, we then re-assessed the strength of the association between each instrumental SNP and its associated gene expression in a particular cell-type using the *F*-statistic[19]. Overall *F-*statistic distributions for each cell-type in Supplementary Figure 4 (IV-gene *F-*statistic range 16.9 - 233, median 29).

Following the above steps only a single SNP was retained as the selected IV for most (96.9%) gene/cell-type/outcome combinations. Less commonly encountered was the occurrence of >1 IV for a particular gene/cell-type/outcome combination. For example, colocalization between a genome-wide significant chromosomal region on 5q35.3 for AD and *cis-*eQTLs for *RASGEF1C* in microglia identified 2,184 SNPs in the colocalised chromosomal region (PP.H4=0.87). Removal of SNPs with a *cis-*eQTL FDR >5% followed by removal of SNPs in LD (r^2^>0.01) with the lead *cis-*eQTL eSNP resolved two independent IVs for *RASGEF1C* in microglia, namely: rs76792388 and rs10077711, with study-wide *cis-*eQTL FDRs of 2.40×10^−4^ and 4.62×10^−2^ respectively. In line with the MR assumptions, we considered each IV to independently instrument *RASGEF1C* expression and both IVs were combined in a single inverse-variance weighted (IVW) MR test to estimate the overall contribution of *cis-* regulatory control of *RASGEF1C* expression to AD risk (MR analysis detailed below).

In total, we identified 167 unique IV SNPs which, because a single IV may instrument the same gene across multiple cell-types and/or co-localise with multiple health outcomes, represented 262 IV-gene/cell-type/outcome combinations. Identifying the causal mechanism by which IVs instrument gene expression is challenging due to the multiple mechanisms by which genetic variants can have an effect on gene expression such as alteration of RNA splicing, disruption of *cis-*regulatory enhancers or promoters etc as well as cell-type specific effects on gene regulation which are poorly annotated[20]. Moreover, from a drug target discovery perspective, the precise mechanism by which an IV influences a gene’s expression is less important for MR than the reliability of the association. Nevertheless, an understanding of the mechanisms of *cis-*regulation can add support to the SNP-gene association. We therefore assessed the IVs first using a cell-type agnostic repository of regulatory variants (SNP2TFBS) affecting predicted transcription factor binding sites[21]. This revealed that 41/167 (24.6%) of the selected IVs are predicted to disrupt TF binding affinity (Fig. 3a). We then assessed the regulatory relationship between an IV and its paired gene in a particular cell-type using an external dataset of cell-type specific assay for transposase-accessible chromatin sequencing (ATAC-seq), H3K27ac ChIP-seq,

**Fig. 3.**
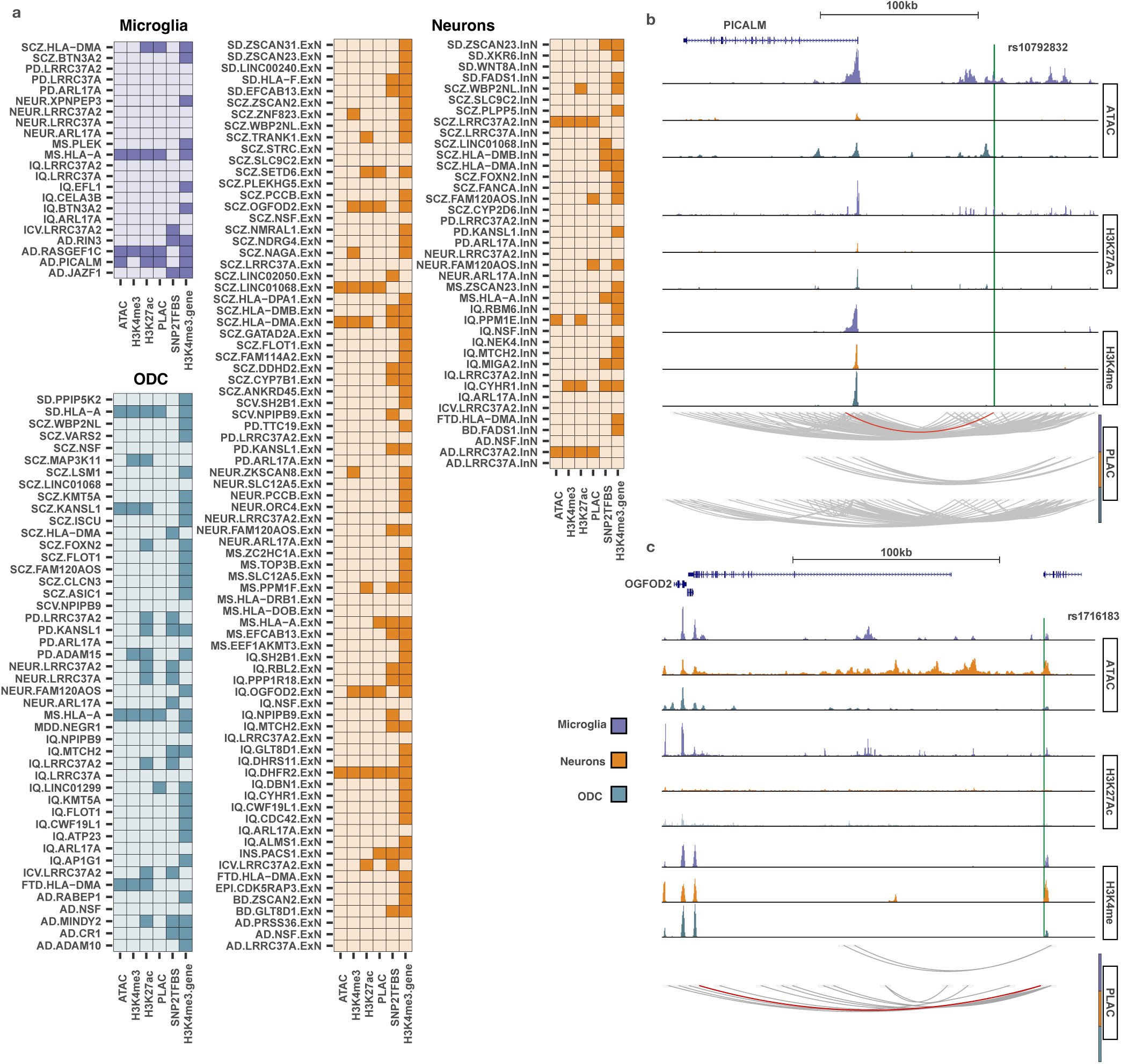
Instrumental variable gene-regulatory landscape. **a**. Cell-type specific gene-regulatory features for the instrumental variables (IVs) in microglia, oligodendrocytes (ODC) and neurons. Each row represents a gene-outcome pair in the indicated cell-type. For neurons, each gene-outcome pair is suffixed with the type of neuron the IV was colocalized in (InN for inhibitory, ExN for excitatory). The first three columns represent (from left to right) the intersection (solid square) between the IV for the indicated gene and an epigenetic feature in that cell-type annotated by ATAC-seq, H3K4me3 ChIP-seq or H3K27ac ChIP-seq. The “PLAC” column indicates whether the IV for the gene in question physically connects to the promoter region of the gene of interest via a PLAC-seq loop in the indicated cell-type. The SNP2TFBS column indicates whether the IV is predicted to disrupt transcription factor binding using the SNP2TFBS database. The H3K4me3.gene column indicates whether the promoter of the gene in question fell within a H3K4me3 ChIP-seq peak in the indicated cell-type. **b**. Genomic map indicating the location of the PICALM instrumental SNP (IV) rs10792832 overlapping a microglial-specific enhancer and connected to the PICALM promoter (red line) via a PLAC-seq loop. **c**. Genomic map indicating the location of the OFGOD2 IV rs1716183 overlapping a neuronal enhancer and connected to the OGFOD2 promoter (red line) via a PLAC-seq loop.

H3K4me3 ChIP-seq and proximity ligation-assisted ChIP-seq (PLAC-seq)[22]. Out of the 186 IV-gene/cell-type/outcome combinations mapping to one or more of the three cell-types for which data were available (neurons, microglia and oligodendrocytes), 40 (21.5%) intersected one or more epigenomic feature supporting the observed cell-type gene regulatory relationship (Fig.3a). For example, for the microglial-specific IV-gene pair rs10792832-*PICALM* (Fig.3b), which colocalises with AD, rs10792832 overlaps a microglial-specific enhancer marked by an H3K27ac peak, is connected to the promoter region of *PICALM* in microglia via a PLAC-seq loop and the PICALM promoter itself overlaps an H3K4me3 peak consistent it with being an active promoter in microglia. For the excitatory neuron-specific IV-gene pair rs1716183-*OGFOD2* (Fig.3c), which colocalises with schizophrenia (SCZ) and intelligence quotient (IQ), rs1716183 overlaps neuronal ATAC and H3K27ac peaks, interacts with the promoter of *OGFOD2* via a PLAC-seq loop in neurons, and the *OGFOD2* promoter overlaps a neuronal H3K4me3 peak.

### Two-sample Mendelian randomisation

For each of the 262 IV-gene/cell-type/outcome combinations we assessed the relationship between the levels of expression of a gene in a particular cell-type with a clinical outcome using the package *MendelianRandomisation*[23]. Here, we used the cell-type specific effect sizes for the IV SNP-gene pair in question as the exposure and the SNP-phenotype effect size from the relevant GWAS as the outcome. In total, we found evidence consistent with a causal interpretation of the association between the levels of expression of a gene and a clinical outcome for 118 genes across 16 brain phenotypes (Summarised in Fig.4a). Of these, 21 genes were inferred to have a causal association to two or more phenotypes (Fig.4b), equating to a total of 149 gene-outcome associations across all phenotypes tested. Whilst there is no single standard by which to benchmark these causal inferences, across all 149 gene-outcome pairs inferred to have a causal association, we find that 132 (88.6%) are reported to have a target-disease association score >0 by the Open Targets Consortium[24] (Fig.4a).

**Fig. 4.**
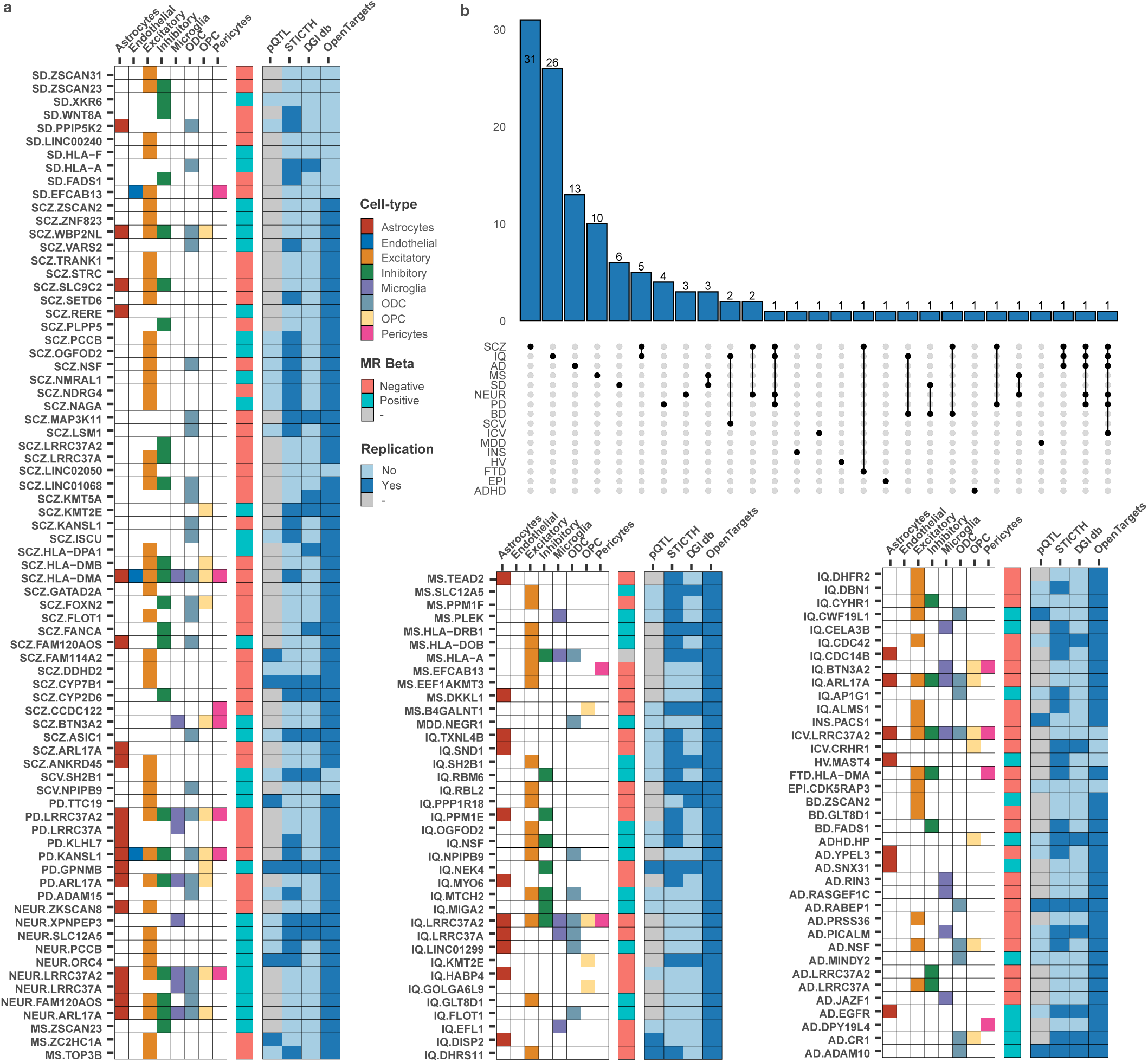
Overview of Mendelian Randomisation results. **a**. Overview of gene-outcome pairs (here labelled with the clinical outcome first to allow causal inferences to be grouped by phenotype) with a significant Mendelian randomisation (MR) association in the indicated cell-type/s. “MR beta” refers to whether the beta coefficients for the IV SNP-gene and SNP-phenotype associations are positively or negatively correlated. A positive correlation can be interpreted as increased gene expression leads to increased outcome risk, whilst a negative correlation can be interpreted as increased gene expression leads to decreased disease risk (or vice versa). Grey squares (MS:*HLA-A*) indicate that the cell-types involved had opposite MR beta directions. Column “pQTL” – solid square indicates that the gene-outcome association at a transcriptional level was reproducible when considering proteins instrumented by the same genetic variant or by a variant in high LD (r^2^>0.8) (grey square indicates that the gene was not assessed in the pQTL dataset due to lack of data). Columns “STITCH” and “DGIdb” – solid square indicates that the protein product for the indicated gene interacts with a known chemical entity from the relevant database. Column “Open Targets” – solid square indicates that the gene-outcome pair have evidence for a target-indication association from the Open Targets Consortium. **b**. Histogram showing the number of genes with an MR-inferred causal association for the indicated phenotype. Genes with an inferred causal association to two or more phenotypes are shown by a solid vertical line connecting the phenotypes. ADHD: attention deficit hyperactivity disorder; EPI: epilepsy; MDD: major depressive disorder; FTD: frontotemporal dementia; HV: hippocampal volume; INS: insomnia; BD: bipolar disease; SCV: subcortical volume (caudate); ICV: intracranial volume; MS: multiple sclerosis; SD: sleep duration; AD: Alzheimer’s disease; NEUR: neuroticism trait; PD: Parkinson’s disease; IQ: full-scale intelligence quotient; SCZ: schizophrenia.

In addition to inferring the causal relationship between genes, cell-types and health outcomes, the present study informs the directionality of the relationships unconfounded by disease-induced changes in gene expression. Knowledge of the directionality of the relationship between the level of expression of a gene and a clinical outcome is critical to informing the therapeutic strategy (i.e., target activation or inactivation), whilst knowledge of the relevant cell-type/s in which they act can inform more precise pre-clinical experimental validations. For example, among the genes inferred to be causal for AD, *PICALM*, encoding phosphatidylinositol binding clathrin assembly protein was first associated with AD in 2009[25]. Currently, no drugs are reported to be in development targeting PICALM as a treatment or prevention strategy for AD[26]. Here, we associate increased *PICALM* expression in microglia with decreased risk of AD (MR *P*=3.03×10^−36^), a finding consistent with the pre-clinical evidence that a reduction in *PICALM* expression increases the development of both amyloid[27] and tau pathologies[28]. Targeting PICALM as a single molecular entity therefore offers the potential to simultaneously modify both amyloid and tau pathologies as a preventative strategy for AD. Notably, of the 16 genes identified by MR in the present study as having a causal association with AD, seven are putatively involved in protein aggregation or trafficking (*PICALM, RABEP1, SNX31, RIN3, PRSS36, NSF* and *MINDY2*), suggesting the absence of drugs in clinical development targeting cellular protein metabolism is a gap in the AD drug development pipeline. Moreover, the MR evidence in support of these genes having a causal association with AD unites the amyloid and tau hypotheses of AD around a single proximal mechanism related to protein trafficking and aggregation.

As a further illustration of the translational value of directionality and cell-type context, we associate increased expression of *GPNMB* (encoding glycoprotein nonmetastatic melanoma protein B) in astrocytes and oligodendrocyte precursor cells (OPCs) with an increased risk of Parkinson’s disease (PD) (MR *P*=3.01×10^−6^ and *P*=1.68×10^−8^ respectively). This directionality of effect was recently independently confirmed by the experimental demonstration that loss of GPNMB activity results in loss of cellular internalization of fibrillar alpha synuclein and reduced pathogenicity, confirming GPNMB inhibition as a candidate therapeutic strategy in PD[29]. Similarly, epidermal growth factor receptor (EGFR) was recently suggested as an AD risk gene following genomic fine mapping based on bulk brain-tissue *cis-*eQTL reference datasets[30]. Here, we explicitly associate decreased EGFR activity in astrocytes with a decreased risk of AD (MR *P*=1.70×10^−7^). This causal inference for *EGFR* is in keeping with EGFR’s known biological relationship to AD, where EGFR inhibition has been shown to ameliorate cognitive dysfunction in different AD models via multiple mechanisms including a reduction in amyloid-beta/tau pathology and inhibition of reactive astrocytes[31]. These findings highlight the potential for EGFR inhibition, including the use of new blood-brain barrier-penetrant EGFR inhibitors[32], as a potential therapeutic strategy in AD.

As well as informing the therapeutic strategy, knowledge of the directionality of an exposure’s effect on an outcome can also inform new biological insights into the causal relationships between genes and phenotypes. For example, five schizophrenia (SCZ) genes (*BTN3A2, FLOT1, KMT2E, OGFOD2, KMT5A*) overlapped with intelligence (IQ) (Fig.4b). For three out of these (*BTN3A2, FLOT1, KMT2E*), the directionality of the gene exposure on SCZ risk and IQ are in the opposite direction (Supplementary Fig.5a). The inverse relationship between SCZ and IQ for these genes may offer an explanation for the clinically observed monotonic relationship between IQ and SCZ – i.e., increasing risk of SCZ with decreasing IQ[33], and therefore targeting them may offer a route to simultaneously alleviating the cognitive deficit associated with SCZ whilst reducing risk of the disease itself. In contrast, where causal genes overlapped between SCZ and neuroticism (*PCCB, FAM120AOS*), the directionality of exposure effect on phenotype was in the same direction (Supplementary Fig.5b). The congruent direction of effect of *PCCB* and *FAM120AOS* on risk of SCZ and neuroticism may partially explain the clinically observed increased risk of SCZ with increasing pre-morbid neuroticism[34], and targeting these genes in SCZ may offer a route to alleviating a maladaptive personality trait associated with SCZ whilst mitigating disease risk. These observations highlight how a phenome-wide approach to single cell-type eQTL-based MR can begin to deconvolute the many complex causal relationships between traits that share overlapping heritability, and thereby improve our understanding of both biology and treatment strategies.

When considering the full set of phenotypes investigated in this study, we observed examples of causal associations across all cell-types of the brain studied (Supplementary Fig.5c), including the lowest abundant cell-types such as pericytes (e.g., multiple sclerosis (MS):*HLA-B*) and endothelial cells (e.g., MS:*ZNHIT6*). Notably, among the 149 gene-outcome combinations inferred to have a causal association, 105 (70.9%) were specific to a single cell-type, suggesting the majority of single gene risk factors for brain outcomes act via a single cell-type as previously observed for immune cell-types and autoimmune disease[35]. Conversely, for clinical outcomes for which multiple risk genes were identified, such as AD, no single cell-type accounted for all the observed heritable effects on phenotypic risk (see Supplementary Fig.6). In situations where the IV-gene pair was inferred to have a causal association with an outcome across more than one cell-type, in all cases the inferred directionality was concordant across the different cell-types.

### Relationship of eQTLs to pQTLs

An implicit assumption in all gene expression studies is that transcript abundance is a valid proxy for protein abundance. A recent comparison of human brain protein QTLs (pQTLs) with eQTLs revealed that a majority pQTLs are also identified as eQTLs[36], although due to lower mapping power for pQTLs not all eQTLs are identified as pQTLs. However, since (currently), proteins represent the dominant category of druggable targets, we assessed the extent to which the association of a clinical outcome with an exposure converges at both the level of transcript and protein abundance. To this end, we used an external dataset consisting of high-throughput mass spectrometry-based protein expression data from bulk-tissue post-mortem brain samples[36]. Across all 118 genes inferred to have a causal association with an outcome in our study, only 51 had a measurable protein expression value in this dataset. Of these 51, 26 had one or more *cis-*pQTL SNP at FDR <5% and of these, 13 (50%) of our MR-inferred causal gene-outcome pairings were reproducible when considering proteins instrumented by either the same genetic variant or by a variant in high LD (r^2^>0.8) (Fig.4a). These results are consistent with the interpretation that for the human brain, causal effects estimated using single-cell snRNA-seq are a valid proxy of a protein’s effect on disease risk. Genes inferred to have a causal association with a clinical outcome at both the level of transcript and protein abundance, and with orthogonal published evidence to support a causal interpretation of the gene-trait association include (trait:gene): PD:*GPNMB*[29], AD:*ADAM10*[37] and AD.*RABEP1*[38].

### Causal genes identified by single cell-type MR identify drug repurposing opportunities

The identification of a causal gene in a specific cell-type is the first step in the development of a new therapy targeting disease risk. To facilitate this, we summarise the cell-type and therapeutic directionality required for risk mitigation for each gene inferred by MR to have a causal association with a brain outcome in Supplementary Fig.6. In contrast to novel drug development, repurposing an existing drug can offer a more rapid route to clinical translation when there is reliable data supporting the target-disease pairing and where the directionality of effect between drug and exposure and between exposure and clinical outcome are known. To explore potential repurposing opportunities, we therefore investigated existing gene-drug interactions using the Drug-gene Interaction Database (DGIdb)[39] and the Sear Tool for Interactions of Chemicals (STITCH)[40]. Of the 118 genes inferred to have a causal association with an outcome, 26 (22.0%) had a reported chemical interaction in DGIdb, and 58 (49.2%) in STITCH (Fig. 4a). These chemical interactions offer a potential tool compound that can be used to experimentally explore the consequences of a drug intervention, or as a starting point for more refined chemistry. Of the drug-gene interactions with a potential for more immediate repurposing, the acid sensing ion channel-1 (encoded by *ASIC1*) was identified by MR as a significant (MR *P*=6.2×10^−4^) risk factor for SCZ associated with increased expression in oligodendrocytes, suggesting that drugs with a negative effect on ASIC1 currents could act to exert a mitigating effect on schizophrenia. Experimentally, over-expression of *ASIC1* has been shown to enhance context fear conditioning in mice and ASIC-like currents have been documented in oligodendrocytes[41]. The licensed potassium sparing diuretic amiloride is a known non-selective blocker of the acid-sensing ion channel-1, currently undergoing evaluation as a prophylactic treatment for migraine (https://clinicaltrials.gov), and highlighted here as a potential novel, non-neuroleptic intervention in schizophrenia.

## DISCUSSION

In this study we mapped genetic effects on gene expression in eight cell-types of the non-diseased control human brain. Single cell-type *cis-*eQTLs were integrated with GWAS loci in a Mendelian randomisation framework to infer causal genes and to identify the cell-types in which they act. In total, we identified 118 genes with MR evidence for a causal association between variation in their levels of expression and susceptibility to one or more brain outcomes. These genes include novel gene-outcome associations as well as genes previously proposed as candidate drug targets for brain disease. For genes with an appropriate measurable pQTL value, we observed a high level of reproducibility of targets identified using cell-type specific gene expression data consistent with causal effects estimated by snRNA-seq being a valid proxy of a protein’s effect on risk.

An important scientific advance of this work is our application of Mendelian randomisation to human brain tissue samples ascertained exclusively from subjects with no history of brain disease. Previous research exploring bi-directional effects between gene expression and disease have suggested that differentially expressed genes are more prone to reveal disease-induced gene expression changes rather than disease-causing ones[9]. Expression QTLs measured in diseased brain samples might therefore be unrepresentative of gene regulatory relationships in the pre-morbid brain. In contrast, the use of non-diseased control human brain samples in a principled Mendelian randomisation framework offers an approach, and a new biological resource, to uncover cell-type specific causal risk factors that are unconfounded by reverse causation and therefore modifiable drug targets for disease prevention.

Our sample size for *cis-*eQTL detection in single cell-types was limited by the substantial difficulties in ascertaining control brain tissue of appropriate quality given the predominant focus of brain banks on brain diseases such as PD, AD, MS etc. Despite these limitations, we report regulatory variants for 10,288 genes across eight cell-types. Future studies that include a larger number of control subjects and an increase in the number of sequenced cells per sample will provide a more granular picture of the role cellular sub-types play in disease aetiology and are likely to lead to additional causal inferences missed by the current study due to sample size limitations. Given the importance of effective target discovery for reducing the costly attrition of drug development in Phase II/III trials, this argues for a concerted global effort to collect control brains in addition to those from people who have died with a neurological or psychiatric diagnosis.

In addition to inferring causality, the present study provides information on the directionality of the association between a gene exposure and a phenotype in a specific cell-type. Knowledge of the direction of effect of an exposure on a health outcome is critical to guiding the directionality of the therapeutic intervention, whilst knowledge of the cell-types via which genes act can aid the design of more precise pre-clinical experiments that may translate better to the human condition. As well as informing therapeutic strategy, knowledge of the directionality of an exposure’s effect on an outcome from MR can also inform new biological insights into the causal relationships between phenotypes when undertaken in a phenome-wide manner as described here. Identification of such shared risk factors across disease categories present opportunities for shared preventative strategies, with a convergence of diverse stakeholders in therapy development hastening drug development. Additionally, as our knowledge of the relationship between existing drug targets and brain disease expands, so too will our ability to predict long-term adverse health effects from candidate therapeutic interventions.

In conclusion, we report a generalizable framework for the selection of genetic instruments and principled conduct of single-cell Mendelian randomisation regardless of starting tissue. The present study highlights novel mechanistic connections between genes, cell-types and phenotypes, prioritises candidate drug targets in their cellular context and establishes a new resource for the interpretation of GWAS-risk alleles in human brain disease and behaviour.

## ACKNOWLEDGEMENTS

This work was supported by the UKRI MRC (Award No: MR/S02638X/1, Award No: MR/W029790/1) and The Alan Turing Institute under the Engineering and Physical Sciences Research Council grant EP/N510129/1. Brain tissue samples and associated clinical and neuropathological data were supplied by the Parkinson’s UK Brain Bank at Imperial, funded by Parkinson’s UK, a charity registered in England and Wales (258197) and in Scotland (SC037554); the Oxford Brain Bank, supported by the Medical Research Council (MRC), Brains for Dementia Research (BDR) (Alzheimer Society and Alzheimer Research UK), Autistica UK and the NIHR Oxford Biomedical Research Centre; the Edinburgh Brain Bank supported by the MRC; and the Amsterdam Medical Centre Brain Bank. We also acknowledge the support of the Epilepsy Society for AS through the Katy Baggott Foundation and for MT and MJ from the Department of Health’s NIHR Biomedical Research Centres funding scheme. A.N. was supported by the UK Dementia Research Institute, which receives its funding from UK DRI Ltd, funded by the UK MRC, Alzheimer’s Society, and Alzheimer’s Research UK. We are grateful for funding support from Epilepsy Society, UK (SP, RB, JM, SMS). This work would not have been possible without the resources of the above Brain Banks and the people and their families who so generously donated brain tissue.

## METHODS

### Samples

Snap-frozen human brain tissue samples from 60 subjects were obtained from the brain tissue banks with full ethical approvals and appropriate material transfer agreements. We complied with all relevant statutory and ethical regulations approved by the Imperial College research ethics committee regarding the use of human post-mortem tissue samples. At the individual brain banks, post-mortem, fresh tissue samples were snap-frozen in liquid nitrogen vapour for 20 minutes before being stored in −80C freezer long term. Immunohistochemistry was undertaken on all samples using adjacent brain tissue (same block) and assessed for beta-amyloid, Tau, TDP43, alpha synuclein and p62. All H&E stains were performed by hand. In the selection of control samples we excluded all samples with a pre-mortem history of neurological or psychiatric disease (at any time) and in all cases, there was no evidence of neurodegenerative or other significant disease processes on neuropathological examination.

### Nuclei isolation and single-nuclei RNA-seq

Single-nuclei RNA-seq (snRNA-seq) data was generated at Imperial College on prefrontal cortex and hippocampus samples ascertained from 60 unique subjects. These brain tissue samples were ascertained from the Imperial College, Oxford University, Edinburgh University or Amsterdam Medical Centre brain tissue banks. Nuclei were isolated as previously described[42] except for a slightly extended douncing during the tissue lysis step (see our previous publication for detailed protocol PMID: 34309761)[43]. Additionally, we included snRNA-seq data on temporal and prefrontal cortex control samples from a further unrelated 87 unique subjects from Roche. Details of the Roche control samples and nuclei isolation are as previously described [12]. In all cases, snRNA-seq data was generated using the 10X Single Cell Next GEM Chip targeting a minimum 5,000 nuclei per sample and libraries prepared using the Chromium Single Cell 3′ Library and Gel Bead v3 kit according to manufacturer’s instructions. cDNA libraries were sequenced using the Illumina NovaSeq 6000 system at a minimum sequencing depth of 30,000 paired-end reads per nucleus.

### snRNA-Seq data mapping

The raw sequencing reads in the FASTQ files were used to align to the human GRCh38 genome and quantified gene counts as UMIs using Cell Ranger *count* (version 5.0.1). For snRNA-Seq reads, we counted reads mapping to introns as well as exons by *--include-introns* option in Cell Ranger (version 5.0.1). As shown in the earlier studies, this results in a greater number of genes detected per nucleus, as well as better cell type classification[44], [45].To build the latest reference genome for read mapping, we followed the recommended building steps by 10X Genomics. We then modified sequence headers in the Ensembl FASTA file, removed version suffix in the Gencode GTF file, defined string patterns for GTF tags, constructed the gene IDs, and filtered the GTF file based on the gene IDs. Finally, the reference genome was created using Cell Ranger *mkref* (version 5.0.1) with default settings[46].

### Genotyping

Donor DNA from samples processed at Imperial College was genotyped using the Illumina Infinium Global Screening Array v2.0. The tool PLINK (version 1.90b6.18) was applied to call genotypes using the default settings[47]. Roche control subject were genotyped as previously described[12]. These genotyped data were harmonized to the hg38 reference genome using bcftools (version 1.9) with the *fixref* plugin (-m flip option)[48], [49]. Prior to imputation, no missing data threshold or minor allele frequency (MAF) or Hardy-Weinberg equilibrium (HWE) filters were applied. Imputation was done on the Michigan Imputation Server (version 1.6.3) using Haplotype Reference Consortium (version r1.1) reference panel of European population[50], [51] with a pre-phasing using Eagle (version 2.4)[52] and imputation using Minimac4[50]. Only bi-allelic SNPs where imputation score (r^2^) was >0.8 were kept. Imperial and Roche samples were merged as previously described [12]. Genetic variants common to imputed genotypes and whole genome sequencing were identified and merged by bcftools (version 1.9)[48]. Post-merging, SNPs with MAF <5% and *P* <10^−6^ in HWE were excluded. Finally, we performed kinship analysis and excluded all samples with a kinship coefficient above 0.2. Following these steps, we retained ∼5.17 million high-quality SNPs in 128 individuals for further analysis.

### Demultiplexing

Sample pools were demultiplexed based on their genotype using the Demuxlet algorithm with the default settings, as previously described[53], [54]. The variable SNPs between the pooled individuals were used to determine which cell belongs to which individual and to identify doublets. Droplets called doublet by Demuxlet were removed from downstream analyses.

### QC and processing of snRNA-Seq data

The quality of snRNA-Seq datasets was assessed using the following metrics: number of total reads per library, sequencing saturation (fraction of reads originating from an already-observed UMI as reported by Cell Ranger *count*), estimated total recovered nuclei, mean of reads per nucleus, number of genes detected, median UMI Counts per nucleus and reads mapped to genome. While the quality of each cell was assessed using filtered feature-barcode matrices (generated using Cell Ranger workflow and *EmptyDrops* implemented in Cell Ranger, version 5.0.1)[12]. For each sample pool, the data was saved as Seurat object by *CreateSeuratObject* function in Seurat (version 4.0.1)[55]. Nuclei exhibiting mitochondrial read proportions higher than 5% and genes expressed in less than 5 nuclei were removed from further analysis. Dimensionality reduction and clustering were conducted based on Seurat’s built-in functions using standard workflow. After clustering, we predicted potential doublets using DoubletFinder (version 2.0.3) based on the filtered matrix, with the assuming doublet formation rate equal to 0.07 as previously illustrated[43], [56]. Potential doublets identified by DoubletFinder were removed. To integrate the samples, we employed the recommended integration method within Seurat using reciprocal PCA (“RPCA”) with default settings. Samples with less than 500 cells were excluded from downstream analysis. Cell-types were assigned using canonical cell-type markers. Specifically, Excitatory Neurons: *SLC17A7, SATB2, VIP, LAMP5*, Inhibitory Neurons: *GAD1, GAD2, SOX6, PVALB*; Astrocytes: *AQP4, GJB6, FGFR3*; Microglia: *CTSS, C1QB, CSF1R*; Oligodendrocyte Precursor Cells (OPC): *CSPG4, PDGFRA, VCAN*; Oligodendrocytes: *MAG, MOG*; Pericytes: *PTGDS, ATP1A2, ITIH5, FLT1, DCN, PDGFRB*; Endothelial Cells: *ACTA2, KCNJ78, ZEB1*.

### eQTL mapping

Raw count matrices were extracted for each cell type, after which counts for all cells were summed per individual, to obtain a single aggregated count value per cell-type per individual. For an individual to be included in the pseudobulk dataset, a minimum of 20 cells in that cell type was required. The aggregated count matrices were then normalised with the *cpm* function (counts per million) from the edgeR package[57] and log-transformed. Mapping of *cis*-eQTLs was performed using MatrixEQTL[11] with a *cis* window of 2Mb (1Mb from each end of the gene) and default parameters. For each cell type, the input consisted of the pseudobulk matrix, genotype matrix, SNP locations file, gene locations file and a covariate matrix including individual-level information for age, sex, post-mortem index (PMI) and sample source. In addition, for each filtered expression matrix, we included the first 40 principal components (PCs) of gene expression as fixed covariates to increase power to detect signals, as previously suggested [12]. We included all genes expressed in at least 3 individuals per cell-type, and genetic variants with at least two individuals in 2 out of the 3 genotypic categories. False Discovery Rate (FDR) using the Benjamini–Hochberg method for both discovery sets was applied [13].

### Validation of eQTLs using a bulk dataset

We obtained the full *cis*-eQTL associations from a recent bulk eQTL dataset (“Metabrain” dataset) performed on 6,518 individuals[14]. To calculate the percentage overlap, we first identified *cis*-eQTLs (SNP-gene pairs) with a study-wide FDR <5% FDR in each cell-type. This identified a total 39,840 SNP-gene pairs for astrocytes, 4,339 for endothelial cells, 140,053 for excitatory neurons, 40,463 for inhibitory neurons, 20,180 for microglia, 66,114 for oligodendrocytes, 21,418 for oligodendrocyte precursor cells and 7,884 for pericytes. The percentage overlap with the external *cis*-eQTL dataset in Metabrain was then calculated based on the total number of SNP-gene pairs also in Metabrain at FDR <5%, divided by the aforementioned numbers.

### Colocalisation analysis

We employed COLOC[16] to perform colocalisation analysis. Briefly, *cis-*eQTLs were generated for each cell type as described above. To prepare the summary statistics for colocalisation analysis, we first performed a liftover from hg19 to hg38 using the *liftOver* function from the *rtracklayer* package[58] and the latest liftover chain file from UCSC (hg19ToHg38.over.chain). For each GWAS trait analysed, the regions were selected based on variants with the most significant genome-wide association in a non-overlapping fashion (meaning each selected region could have more than one genome-wide significant SNP). The summary statistics were scanned using the *ld_clump* function of the *ieugwasr* package[59] and only the top genetic variant in a window of 1Mb was kept (500kb on either side of the variant). The regions were then re-populated with the full list of variants situated within the 1Mb window of each region to then be used in the colocalisation analysis. To perform single-cell eQTL colocalisation, the full *cis*-eQTL associations for each cell type were intersected with variants in each GWAS trait on a per-region basis. For each region, COLOC was then used iteratively in a binary fashion between the GWAS and all cell-type/gene combinations using default priors. Each cell type/gene combination was considered as a single trait (such as astrocyte/*APIP*), i.e., the total number of colocalisation tests performed would be equal to the number of genes multiplied by the number of cell-types. For example, in a region with 20 genes, a total of 160 when considering 8 cell types. This was repeated for every region of genome-wide significance in each GWAS. For downstream analysis, traits with a regional posterior probability (PP.H4) above 0.5 were retained. For quantitative GWAS traits, `type=“quant”` was specified in the COLOC input. For case/control GWAS traits, `type=“cc”` was specified in the COLOC input. For all *cis*-eQTL traits, `type=“quant”` was specified in the COLOC input.

### Mendelian randomisation

Mendelian randomisation was performed using the *MendelianRandomisation R* package[23]. For each GWAS, regions around colocalised traits (cell/gene combination) with a posterior probability (PP.H4) of more than 0.5 were carried forward to MR. The genetic variants were then filtered to satisfy the mendelian randomisation assumptions. First, to ensure the robustness of our instrumental variables, we only kept variants in that region with an association with the gene at FDR below 5%. Following this, we excluded all variants in high LD (r^2^ >0.01) with the lead variant(s). In the large majority of cases (>90%), only one instrumental variable (IV) was retained. Then, we applied Mendelian randomisation using the *mr_allmethods* function specifiying `“ivw”` (with a fixed-effects meta-analysis for more than one IV and the ratio method when there was only one IV) as the method to be used, using the cell-type specific effect sizes for the gene in question as the exposure and the GWAS effect size as the outcome.

### Intersection with protein-QTL dataset

To assess whether our MR hits had actionable potential evidenced by protein expression, we sought to identify overlaps with published pQTL datasets. We obtained two published pQTL summary stats from a study recently conducted using samples from the dorsolateral prefrontal cortex[36]. The first contained all individuals in the study, which included samples with Alzheimer’s Disease, while the latter only contained samples obtained from individuals with no cognitive impairment (NCI). To perform our overlap, we first intersected exact SNP-gene pairs obtained from our MR results (instrumental variable(s)-gene). In addition, we extended this overlap for SNPs in high LD (r^2^ >0.8) with the instrumental variable(s). This was assessed using the *LDproxy* function from the *LDLinkR* package[60], specifying `“CEU”` as the population to be used.

### Intersection with epigenetic data

To assess the cis-regulatory evidence of our MR hits, we intersected our hits with data from a recently published article on cell-type specific epigenetic regulation assessed through Histone ChIP-seq and PLAC-seq[22]. We obtained the processed and filtered *bed* files from the author’s GitHub page (https://github.com/nottalexi/brain-cell-type-peak-files). For our IV intersection, we first performed a liftover from hg38 to h19, as the peak files were on this build, before the intersection. For the gene promoter intersection, we first obtained gene promoters from the *TxDb*.*Hsapiens*.*UCSC*.*hg19*.*knownGene* package using the *promoters* function from the *GenomicRanges* package[61], specifying a maximum range of 5,001 bp (to ensure overlap with the PLAC-seq fragments, which are 5kb long).

### Intersection with drug targets

We investigated whether our MR targets were potentially actionable through therapeutic targeting based on available protein interaction databases. To do so, we downloaded the following interaction databases; DGIdb[39] STITCH[40], and OpenTargets[24]. For STITCH, we downloaded the protein/chemical links dataset and kept all connections with a “combined score” of 0.9 and above (which is equivalent to the highest confidence of connections according to the STITCH guidelines), obtained from http://stitch.embl.de/. We converted protein ENSEMBL IDs using the *biomaRt* package[62]. For DGIdb, we downloaded the latest set of interactions (“interactions.tsv”, “genes.tsv” and “drugs.tsv” of February 2022), obtained from https://www.dgidb.org/downloads. For OpenTargets, all data was downloaded from https://platform.opentargets.org/downloads. We performed two sets of analysis. First, we tested whether the MR genes in question were also putative targets for the trait analysed in OpenTargets. To do so, we downloaded the “Associations – direct (overall score)” dataset, which contains scores for putatively important risk genes. For our analysis, we intersected all genes with a score above 0. Secondly, we tested whether our targets had been previously used for therapeutic design. Hence, we downloaded the “Target” and “Drug” datasets to assess whether this was the case and matched these to our MR genes.

### Processing of GWAS summary statistics

We standardised all GWAS studies to contain the following headers; “chr” for chromosome position, “pos” for a base-pair position, “rsid” for SNP id, “pval” for association p-value, “b” for the effect size, “se” for the standard error, “A1” for the effect allele, “A2” for the other allele and “MAF” for the minor allele frequency. In cases where the effect size was missing but the Z-score was available, we calculated the effect size (beta regression coefficient) and standard error using a previously described formula[63]. When the Odds Ratio (OR) was included but not the effect size, we did a natural logarithmic conversion to obtain the effect size.

## Data availability

The datasets generated during and/or analysed during the current study will be made available at the point of publication deposited within the European Genome-phenome Archive.

## Figures

Most figure panels were generated programmatically in *R* using *ggplot2*[64] with the exception of Fig. 2b-c which were generated using *gassocplot2* (https://github.com/jrs95/gassocplot2). Fig. 1a was created with BioRender.com (full licence). Figure 3b-c was created using the custom tracks on the UCSC genome browser (https://genome.ucsc.edu/) as previously illustrated[22].

---

**Supplementary Fig. 1.**
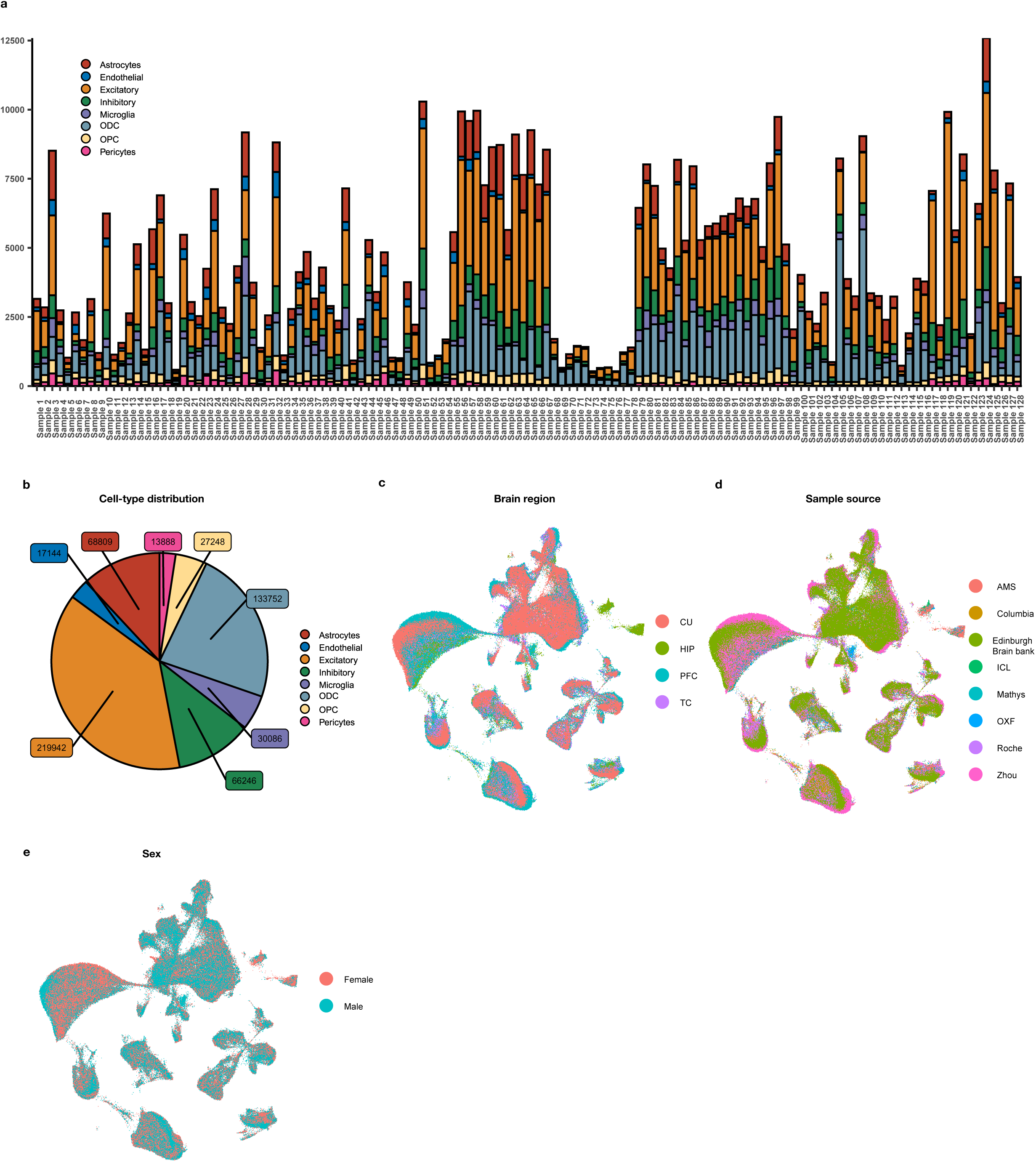
Overview of snRNA-seq on 128 individuals. Following integration, single-cell and sample quality control, we obtained a total of high-quality 577,115 single-cells across 128 individuals. **a.** Number of cells per cell-type sequenced across all individuals used in the study. **b.** Total number of cells discovered across the 8 major brain cell-types. **c.** Distribution of cell-type clusters, annotated by sample source (and/or study). **d.** Distribution of cell-type clusters, annotated by brain region (CU; Cortex (unspecified), HIP; Hippocampus, PFC; Prefrontal cortex, TC; Temporal cortex). **e.** Distribution of cell-type clusters, annotated by sex

**Supplementary Fig. 2.**
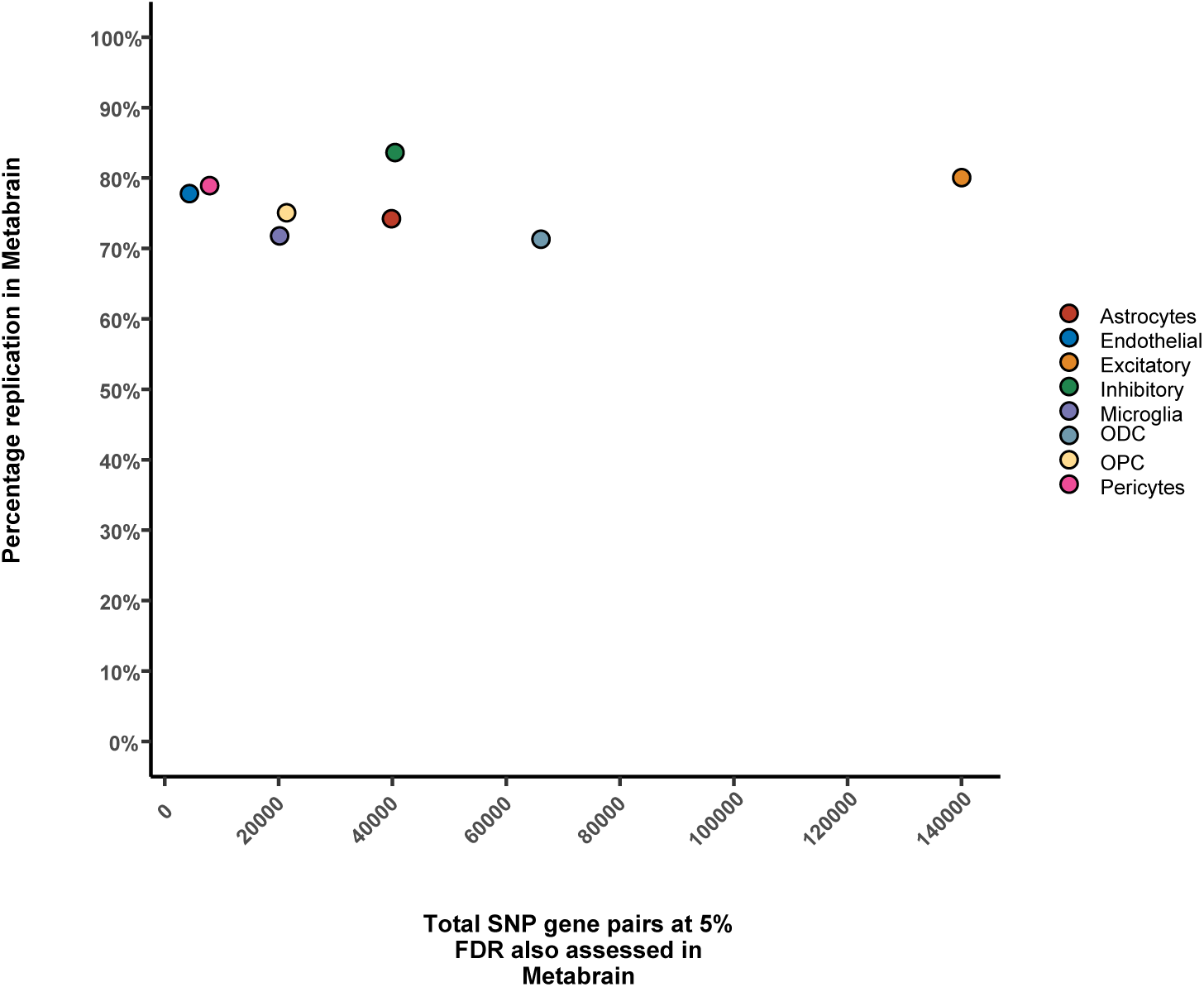
Replication of *cis*-eQTLs in the Metabrain cohort. Our *cis*-eQTL discovery was validated in a large bulk RNA brain dataset (Metabrain) comprising of 6,518 individuals. Each point represents the percentage (*y*-axis) of FDR-significant (<5%) *cis*-eQTLs in a specific cell-type in our cohort that was also of FDR significance (FDR<5%) in the metabrain cohort. The *x*-axis represents the total number of SNP-gene pairs replicated.

**Supplementary Fig. 3.**
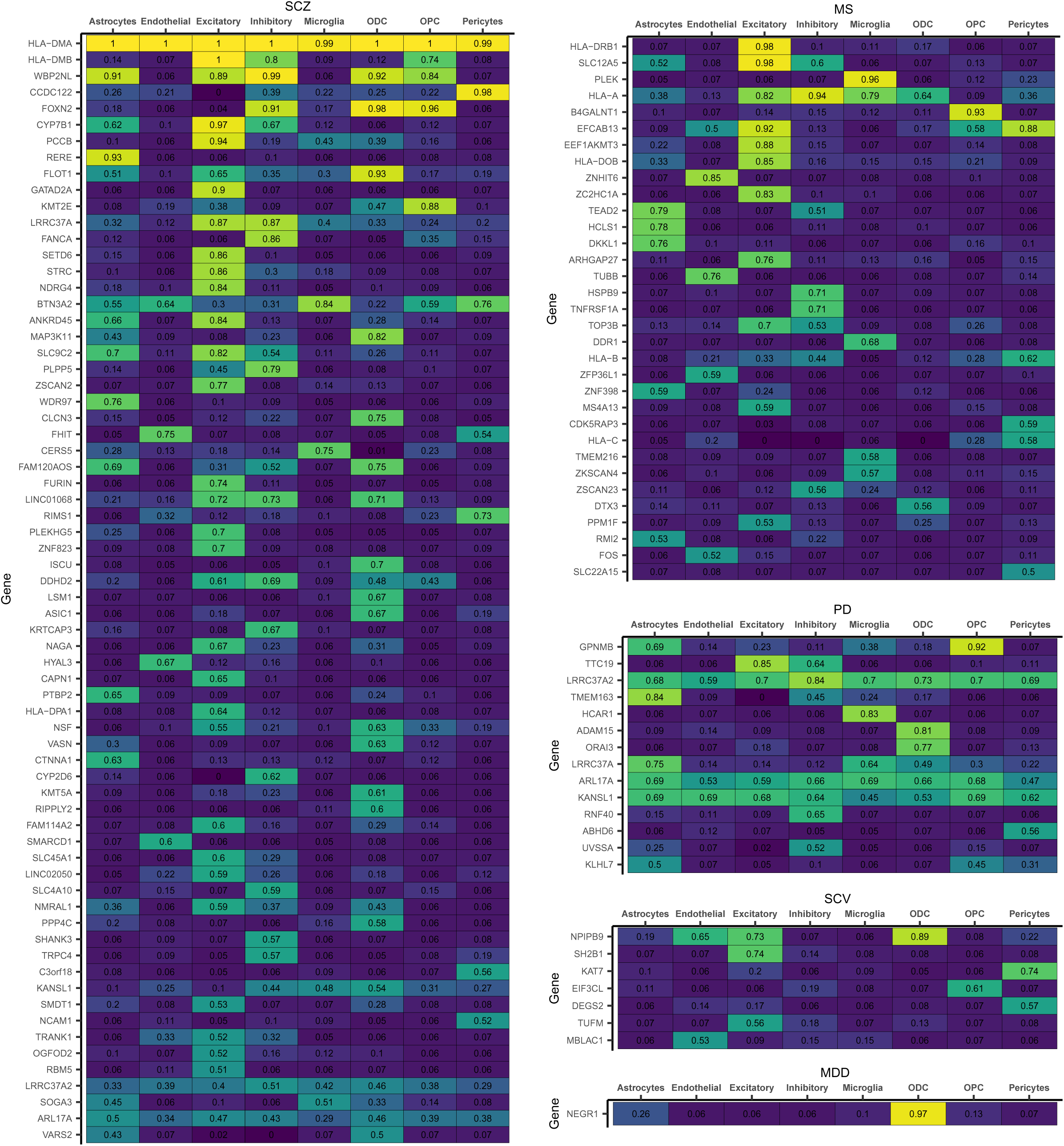
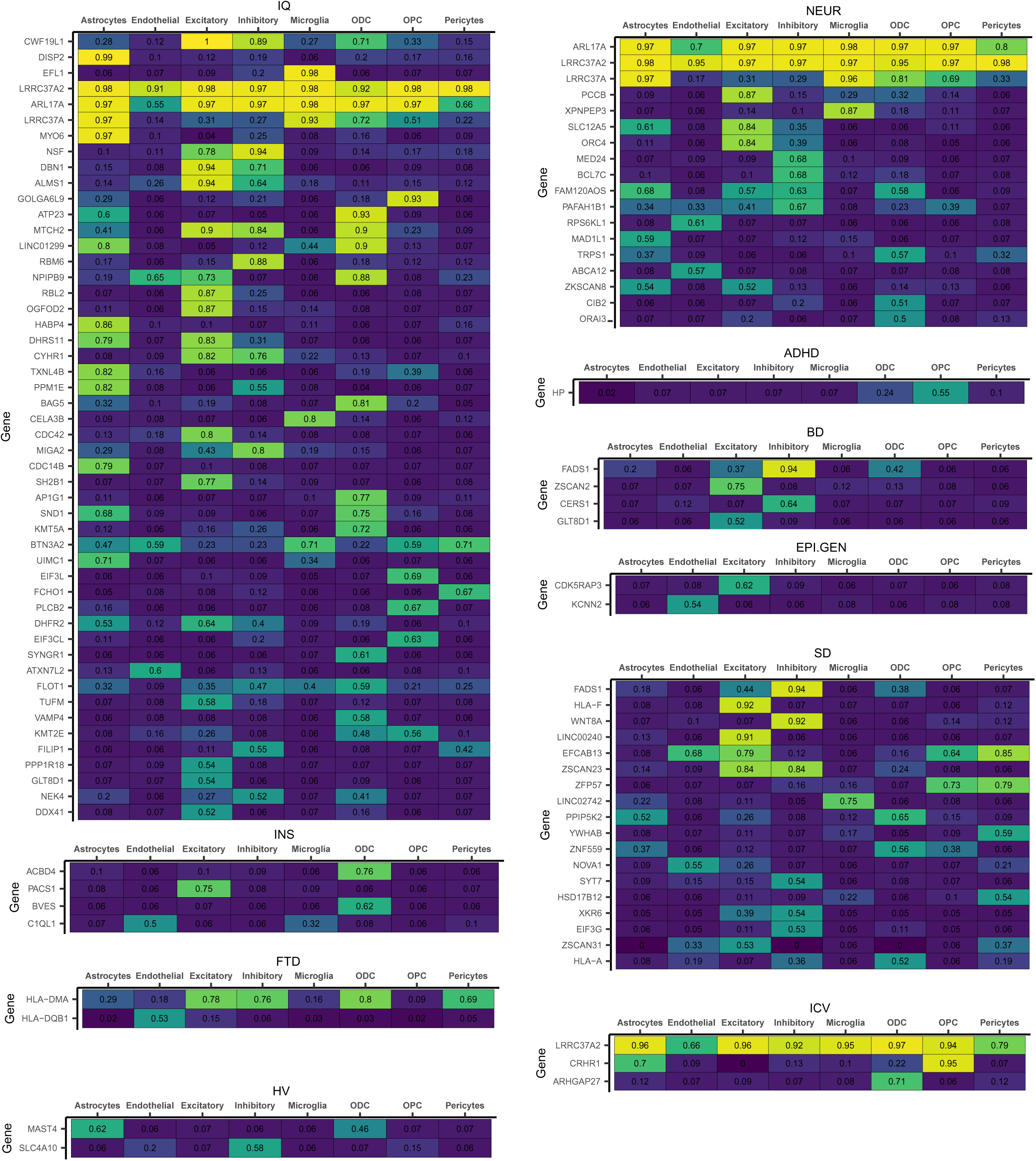
Colocalisation results. Each heatmap shows the posterior probability (PP.H4>0.5) for a shared genetic signal for a SNP-gene (i.e., *cis*-eQTL) pair (row) in a particular cell-type (column) and a genome-wide significant GWAS locus within a given trait (SD: sleep duration; SCZ: schizophrenia; SCV: subcortical volume caudate; PD: Parkinson’s disease; NEUR: neuroticism; MS: multiple sclerosis; MDD: major depressive disorder; IQ: intelligence; INS: insomnia; ICV: intracranial volume; HV: hippocampal volume; FTD: Frontotemporal Dementia; EPI.GEN: genetic generalized epilepsy; BD: bipolar disorder; ADHD: attention deficit hyperactivity disorder, AD: Alzheimer disease).

**Supplementary Fig. 4.**
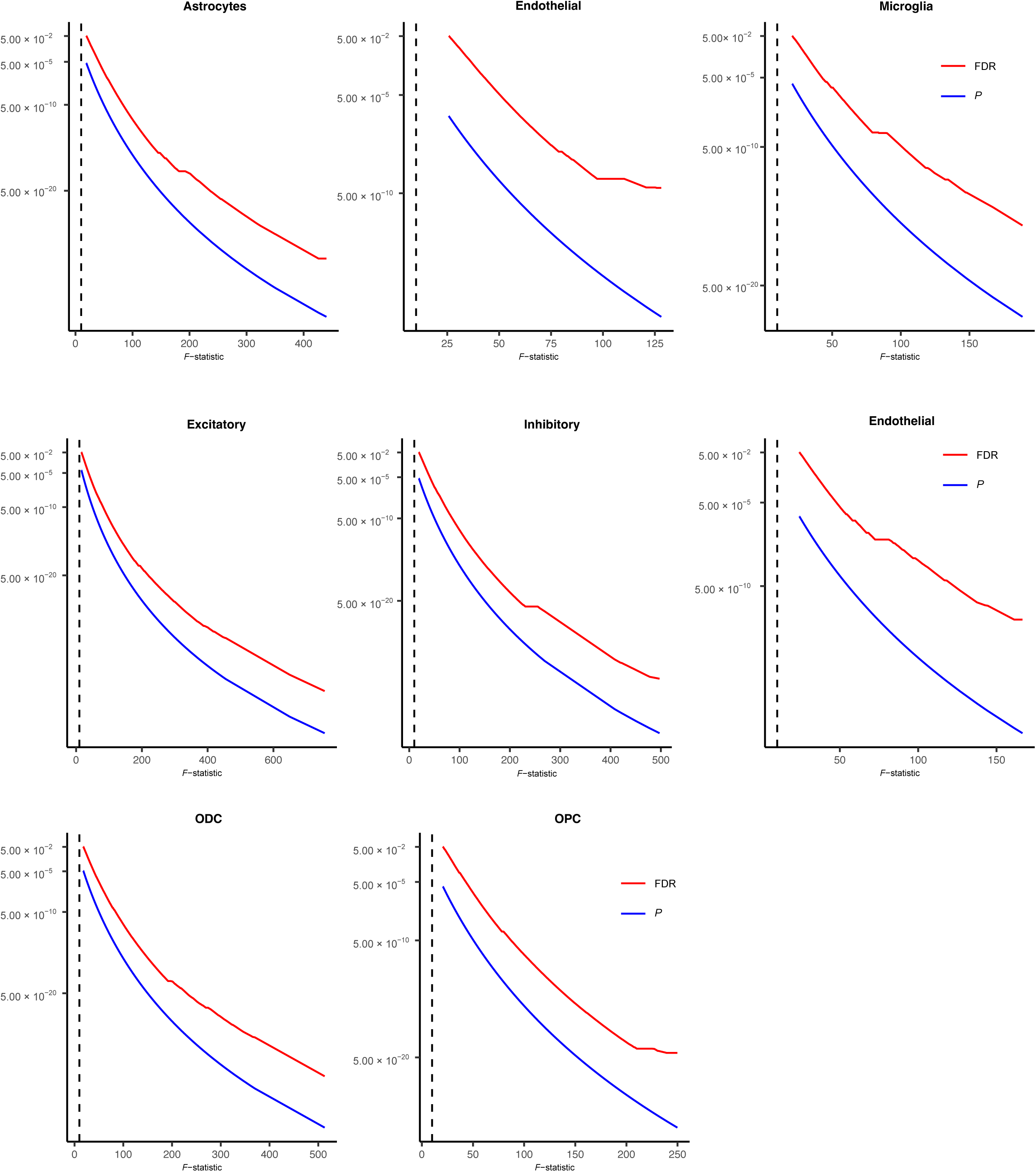
*F*-statistic distributions of *cis*-eQTLs. Mendelian Randomization necessitates selection of robust instrumental variables (*F*-statistic > 10, denoted by the dotted line). In each plot (per cell-type), the *x*-axis represents the *F*-statistic for a given association, and the *y*-axis represents the significance of the association for a given *F*-statistic value (p-value (*P*) in blue, FDR in red).

**Supplementary Fig. 5.**
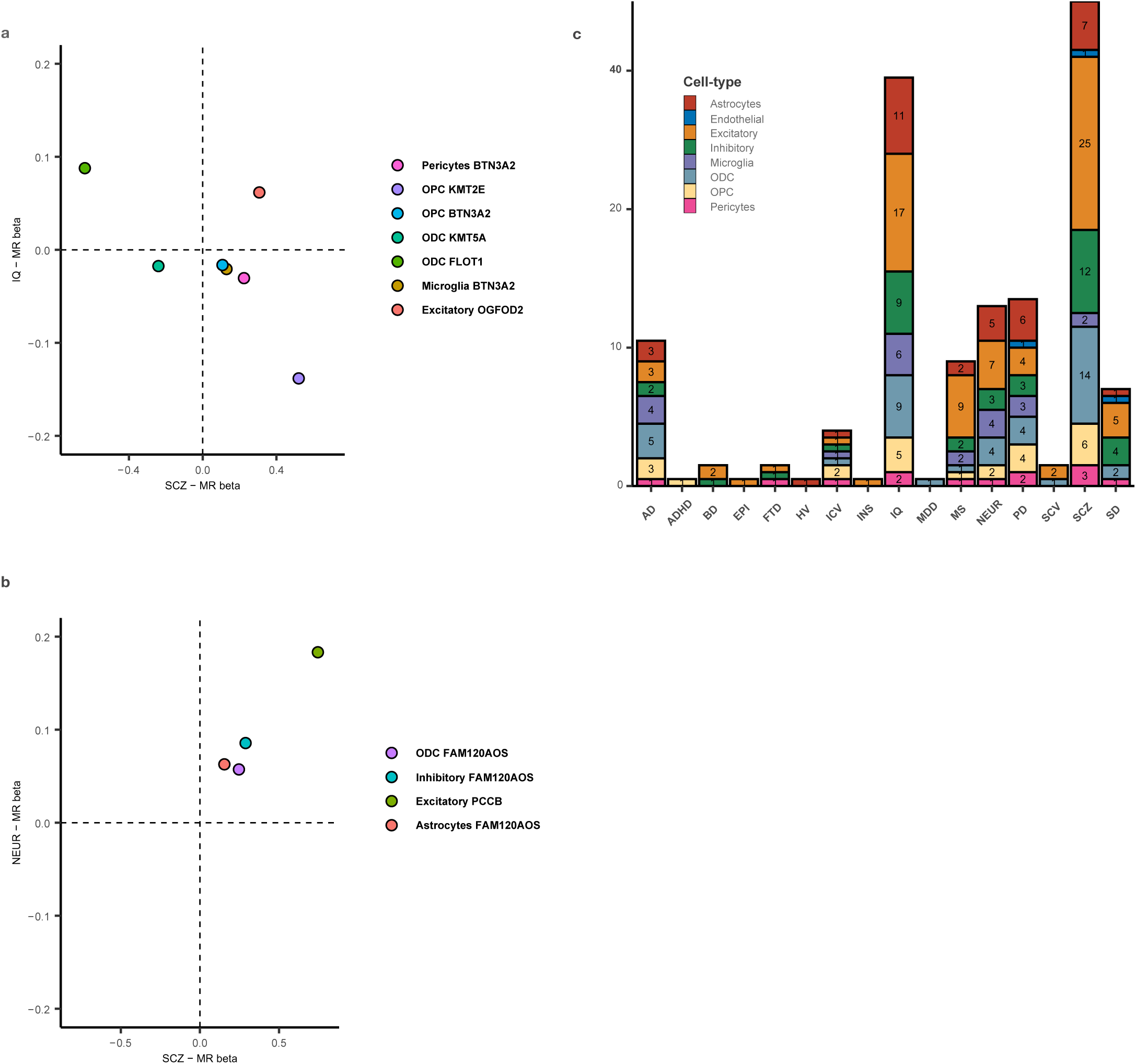
Inferred causal directionality and cell-type specificity. **a**. Cell-type/gene combinations overlapping only between schizophrenia (SCZ) and intelligence quotient (IQ). The *y*-axis represents the MR effect size (beta regression coefficient) for a given cell-type/gene pair in IQ. The *x*-axis represents the MR effect size for that same cell-type/gene pair in SCZ. For example, cell-type/gene pairs in the lower right quadrant (OPC-KMT2E, Pericytes-BTN3A2, Microglia-BTN3A2 and OPC-BTN3A2) indicate a positive MR beta regression coefficient for SCZ but negative for IQ (i.e., increased gene expression for these genes is associated with increased risk of SCZ and reduced IQ). **b**. Cell-type/gene combinations shared only between SCZ and neuroticism (NEUR). The cell-type/gene pairs in the upper right quadrant (Astrocytes-FAM120AOS, Inhibitory-FAM120AOS, ODC-FAM120AOS, Excitatory-PCCB) indicate a positive MR beta regression coefficient for both SCZ and neuroticism (i.e., increased gene expression for these genes is associated with increased risk of both SCZ and neuroticism). **c**. Overview of cell-type proportions per trait for all significant MR results for that trait.

**Supplementary Fig. 6.**
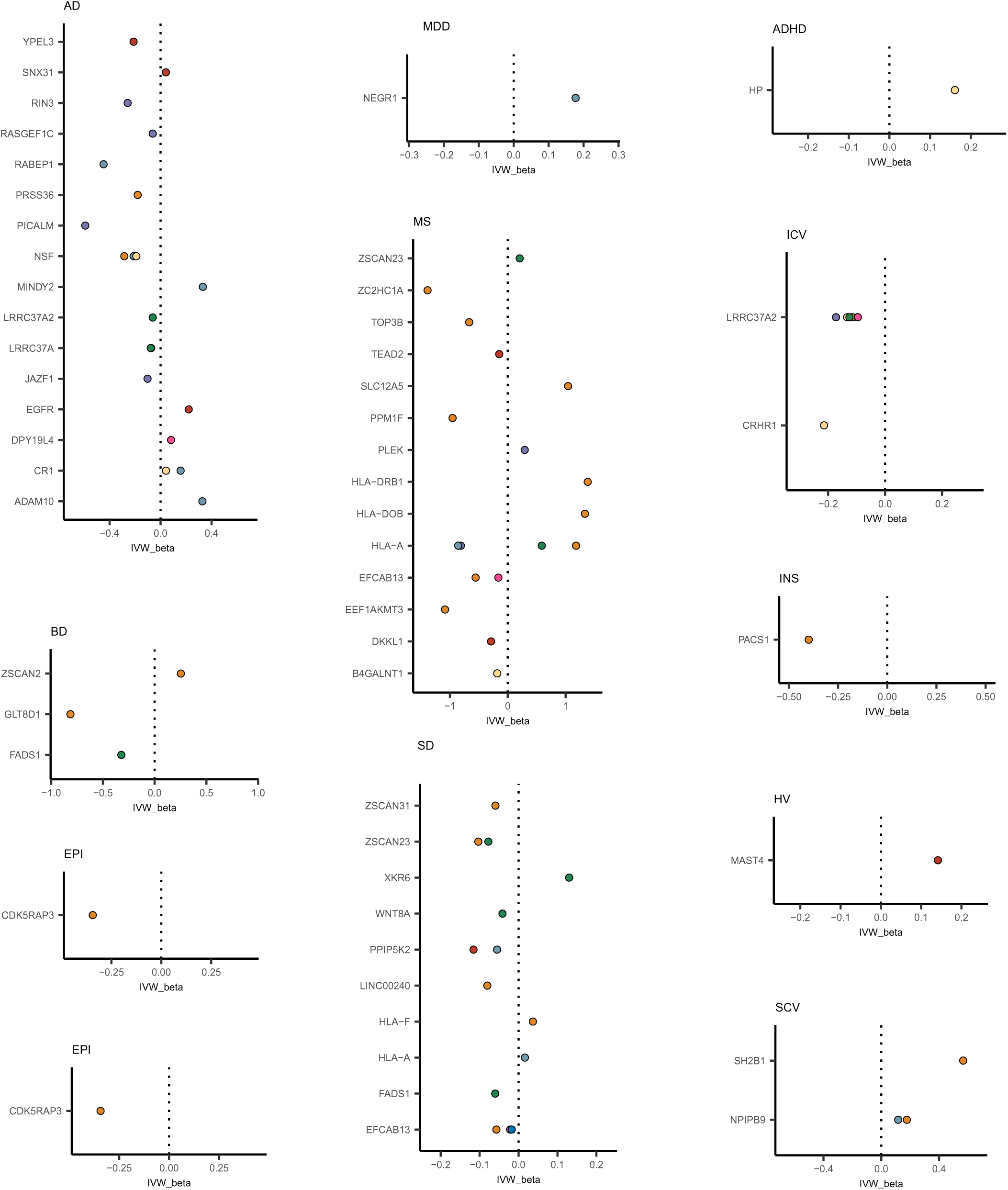
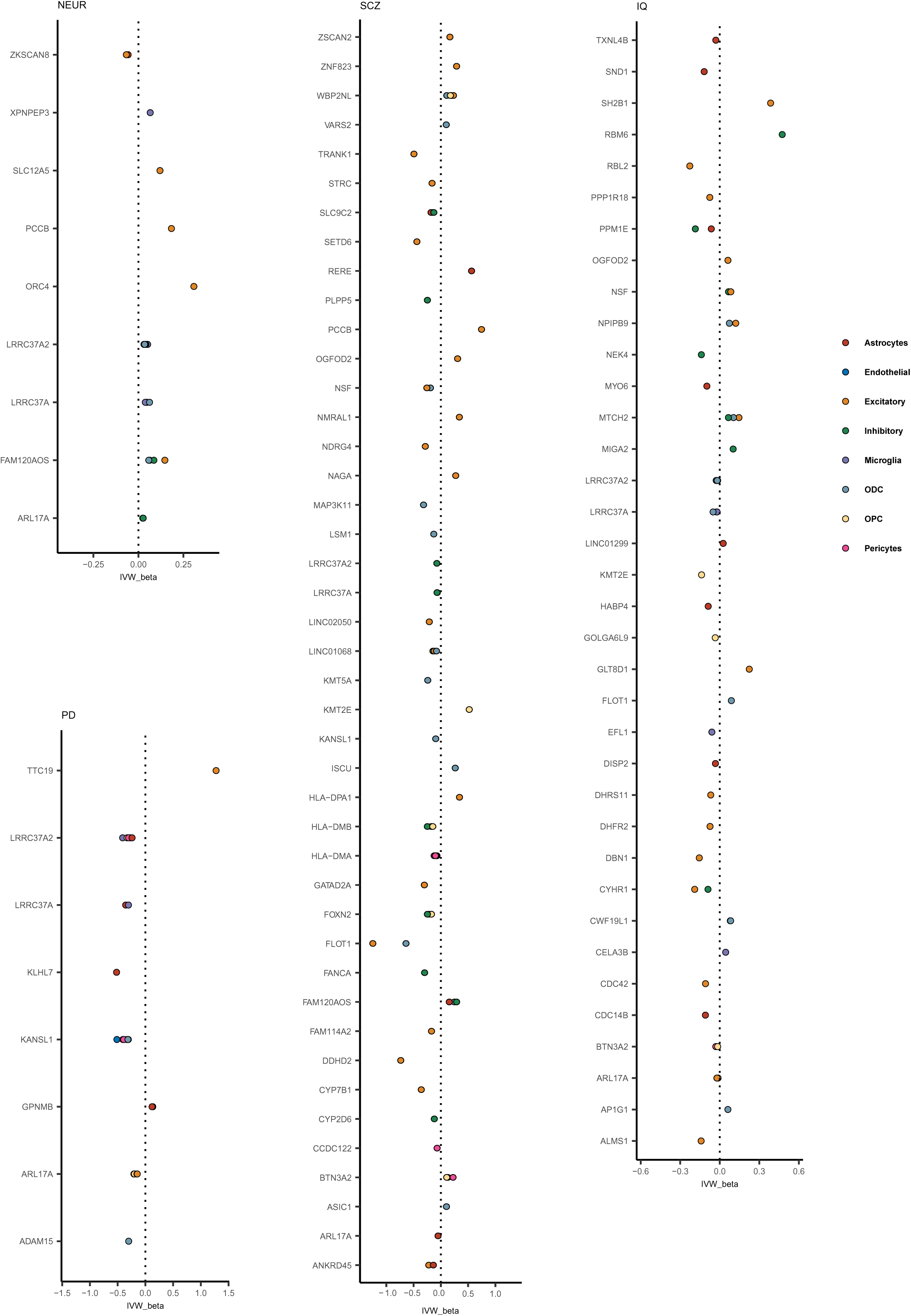
Directionality for all MR hits in across all traits (denoted in each plot title). The *y*-axis denotes the gene for the MR hit, and the *x*-axis denotes the MR beta regression coefficient, indicated by the dot (coloured by cell-type). The dotted line is centered on zero, meaning hits on the right represent a positive beta (increased expression relates to increased risk, requiring target inactivation) whereas on the left represent a negative beta (increased expression relates to decreased risk, requiring target activation). ADHD: attention deficit hyperactivity disorder; EPI: epilepsy; MDD: major depressive disorder; FTD: frontotemporal dementia; HV: hippocampal volume; INS: insomnia; BD: bipolar disease; SCV: subcortical volume (caudate); ICV: intracranial volume; MS: multiple sclerosis; SD: sleep duration; AD: Alzheimer’s disease; NEUR: neuroticism trait; PD: Parkinson’s disease; IQ: full-scale intelligence quotient; SCZ: schizophrenia.

